# Packing the Standard Genetic Code in its box: 3-dimensional late Crick wobble

**DOI:** 10.1101/2021.01.18.427168

**Authors:** Michael Yarus

## Abstract

Minimally-evolved codes are constructed with randomly chosen Standard Genetic Code (SGC) triplets, and completed with completely random triplet assignments. Such “genetic codes” have not evolved, but retain SGC qualities. Retained qualities are inescapable, part of the logic of code evolution. For example, sensitivity of coding to arbitrary assignments, which must be <≈ 10%, is intrinsic. Such sensitivity comes from elementary combinatorial properties of coding, and constrains any SGC evolution hypothesis. Similarly, evolution of last-evolved functions is difficult, due to late kinetic phenomena, likely common across codes. Census of minimally-evolved code assignments shows that shape and size of wobble domains controls packing into a coding table, strongly shifting accuracy of codon assignments. Access to the SGC therefore requires a plausible pathway to limited randomness, avoiding difficult completion while packing a highly ordered, degenerate code into a fixed three-dimensional space. Late Crick wobble in a 3-dimensional genetic code assembled by lateral transfer satisfies these varied, simultaneous requirements. By allowing parallel evolution of SGC domains, it can yield shortened evolution to SGC-level order, and allow the code to arise in smaller populations. It effectively yields full codes. Less obviously, it unifies well-studied sources for order in amino acid coding, including a stereochemical minority of triplet-amino acid associations. Finally, fusion of its intermediates into the definitive SGC is credible, mirroring broadly-accepted later cellular evolution.

## Introduction and approach

The form of the Standard Genetic Code (SGC) offers authoritative information about its origin. By calculating evolved coding tables (Yarus 2021b), the SGC’s implications can be investigated. Comparing coding tables evolved via different pathways, more frequent SGC-like results quantitatively signal superior explanations. Consequently, initial hypotheses about code descent can be improved. In fact, respecting Bayes’ theorem, multiple successful explanations rapidly strengthen an accurate hypothesis by Bayesian convergence (Yarus et al. 2005).

The existing result is late Crick wobble (Yarus 2021a). “Late” implies that wobble was deferred, being preceded by unique triplet pairing assignments. Unique base pairing does not require support from a highly evolved allosteric ribosome (Moazed and Noller 1990; Ogle et al. 2001), or a specific, highly optimized tRNA anticodon loop-and-stem structure (Yarus 1982; Uhlenbeck and Schrader 2018), or control of isomerization in wobble-paired bases (Westhof et al. 2019). Accurate Crick wobble (Crick 1966) would therefore likely be a later, more modern code refinement. Such late wobble also shows superior ability to fill an SGC-like coding table, and offers more probable SGC access to evolving codes (Yarus 2021b, 2021a). Moreover, the first wobble (the SGC necessarily uses ambiguous coding) probably resembled simplified Crick wobble, defined here as translation of NNY (Y = pyrimidine) and NNR (R = purine) codons, each with one adaptor RNA (Crick 1966). For comparison, superwobble, translation of four NNY/R codons with one unmodified adaptor (Alkatib et al. 2012), less frequently yields the SGC (Yarus 2021a). Other evolutionary obstacles to SGC-like coding can be defined similarly, within this simulated framework.

## Results

### Random, minimally-evolved codes

Minimally-evolved codes are random coding tables, or differ in only defined ways from randomly-assigned coding tables (Methods). They have determined fractions of randomly-chosen SGC codons, completed with completely random assignments. A property appearing in such a minimally-evolved state is likely intrinsic to code evolution. This construction defines crucial evolutionary problems for the SGC, and allows evaluation of claimed solutions.

### Inappropriate assignments

The simplest among the inevitable is overall sensitivity of code evolution to nonspecific triplet assignment. Fig. 1A plots assignment accuracy; the probability of codes with 0, ≤ 1, ≤ 2, ≤ 3 or ≤ 4 mis-assignments (abbreviated “mis”) relative to the SGC. Code accuracies are shown versus the fraction (Prand) of random triplet meanings (20 amino acids / termination / initiation) rather than canonical SGC triplet assignments. Fig. 1A averages 10^4^ independent coding table evolutions, and is thus not tied to any particular choice of SGC or randomized codons. Accuracy is determined after encoding 20 functions, then late Crick wobble (Yarus 2021b, 2021c, 2021a). Fig. 1 results resemble previous calculations of code order (spacing, distance and chemical order taken together) versus Prand. In particular, approaching SGC order or acquiring SGC-like assignments (Fig. 1A) requires that random codon assignment be <≈ 15% of the total, preferably <≈ 10% (Yarus 2021b). SGC-like coding is very sensitive to probability of random assignment (Prand), and sensitivity increases as resemblance to the SGC increases.

**Figure 1A.**
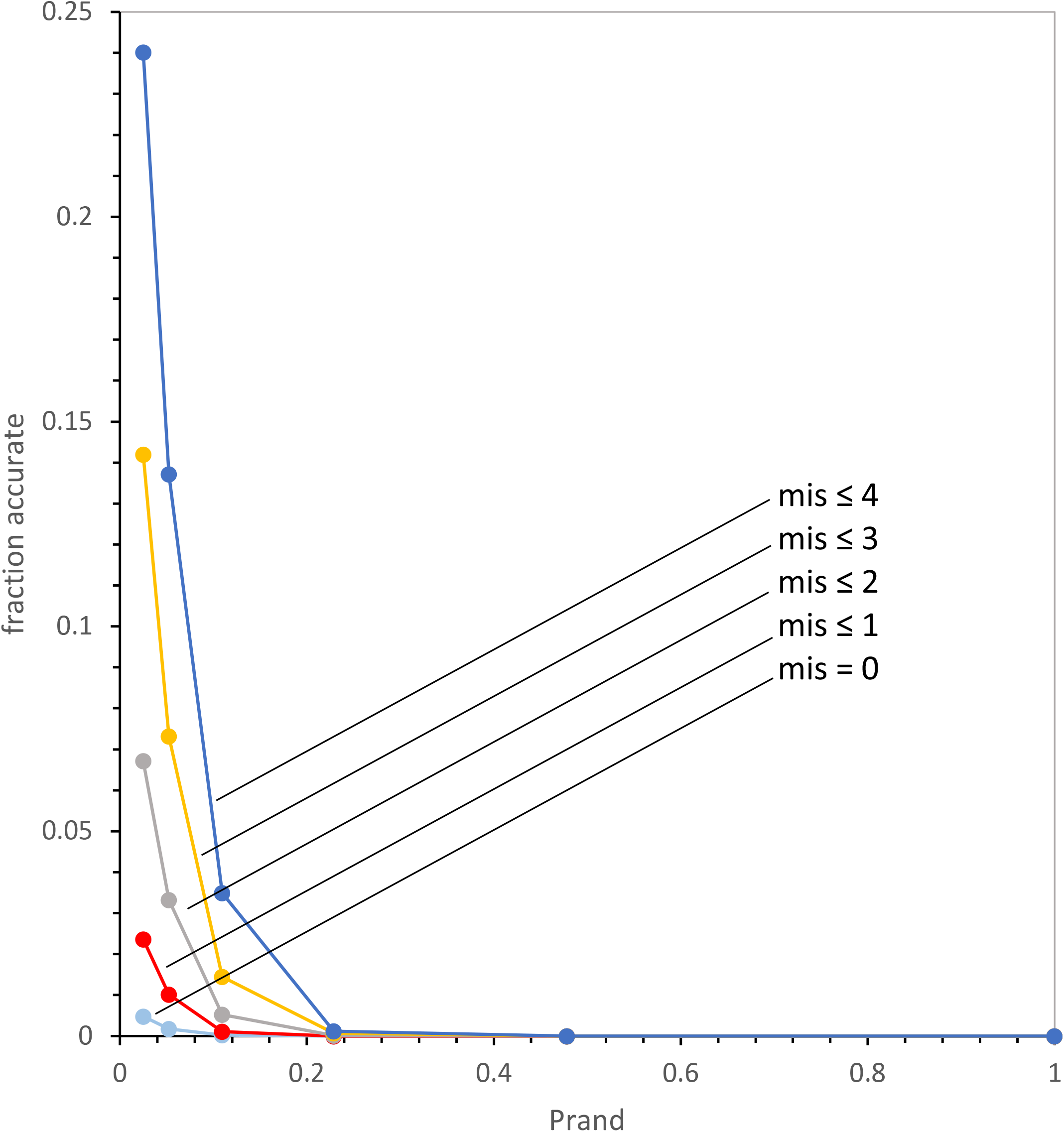
Sensitivity of code evolution to randomness. Fraction accurate = the fraction of 10^4^ evolutions that have indicated accuracy. Accuracy is measured by counting mis-assignments relative to the SGC; mis = 0 (no errors), mis ≤ 1 (single error), mis ≤ 2, mis ≤ 3, mis ≤ 4. Prand = fraction completely random assignment; thus (1 – Prand) = fraction SGC assignments, chosen randomly from an SGC table. Results are for late Crick wobble evolution to 20 encoded functions.

### Inappropriate assignments: a more revealing graphic

These conclusions are fortified by a more informative plot. Fig. 1B posits a minimally-evolved coding table using random assignment probabilities Prand. Predicted code accuracies are approximated in Fig. 1B using the binomial distribution, for a coding table with the same triplet occupancy in Fig. 1A (Methods).

**Figure 1B.**
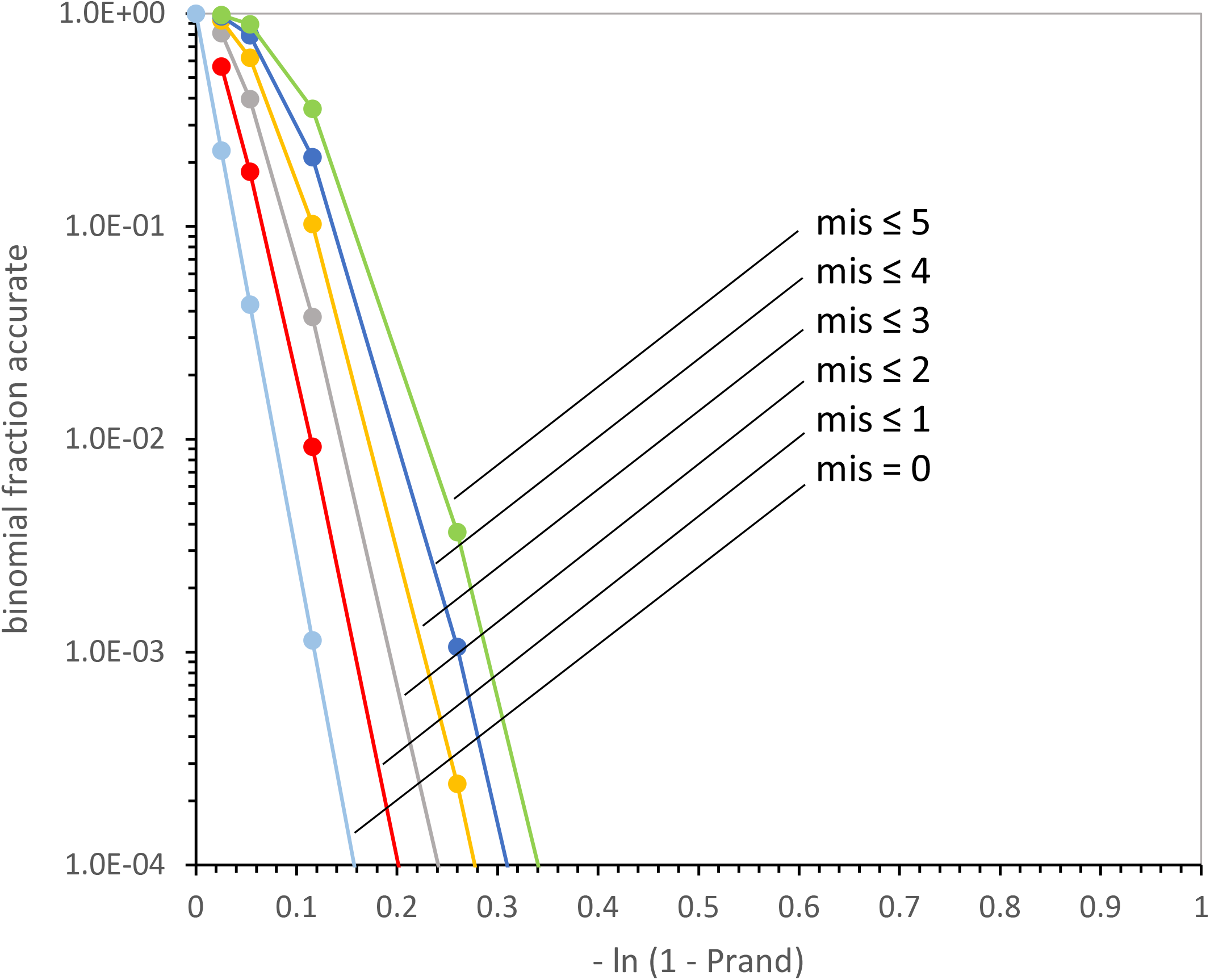
A better portrayal: calculated binomial sensitivity to randomness. Calculated binomially distributed probabilities of choosing 58.6 triplets (the mean in Fig. 1A) with varied Prand (see Methods). Accuracy color coding as in Fig. 1A.

Fig. 1C is data for minimally-evolved codes as in Fig. 1A, but without mutational capture of initial assignments by codons related by single mutational changes. Clearly these minimally-evolved codes, which have late Crick wobble, and all other characteristics of Fig. 1A except capture, greatly resemble the true randomness of Fig. 1B. However, Fig. 1C makes it much clearer that the rarity of accurate codes is innate. As random assignment increases (Prand as x-axis trends similarly to -ln (1 – Prand); Methods), the frequencies of accurate codes, particularly those identical to the SGC (mis = 0), fall exponentially and indefinitely.

**Figure 1C.**
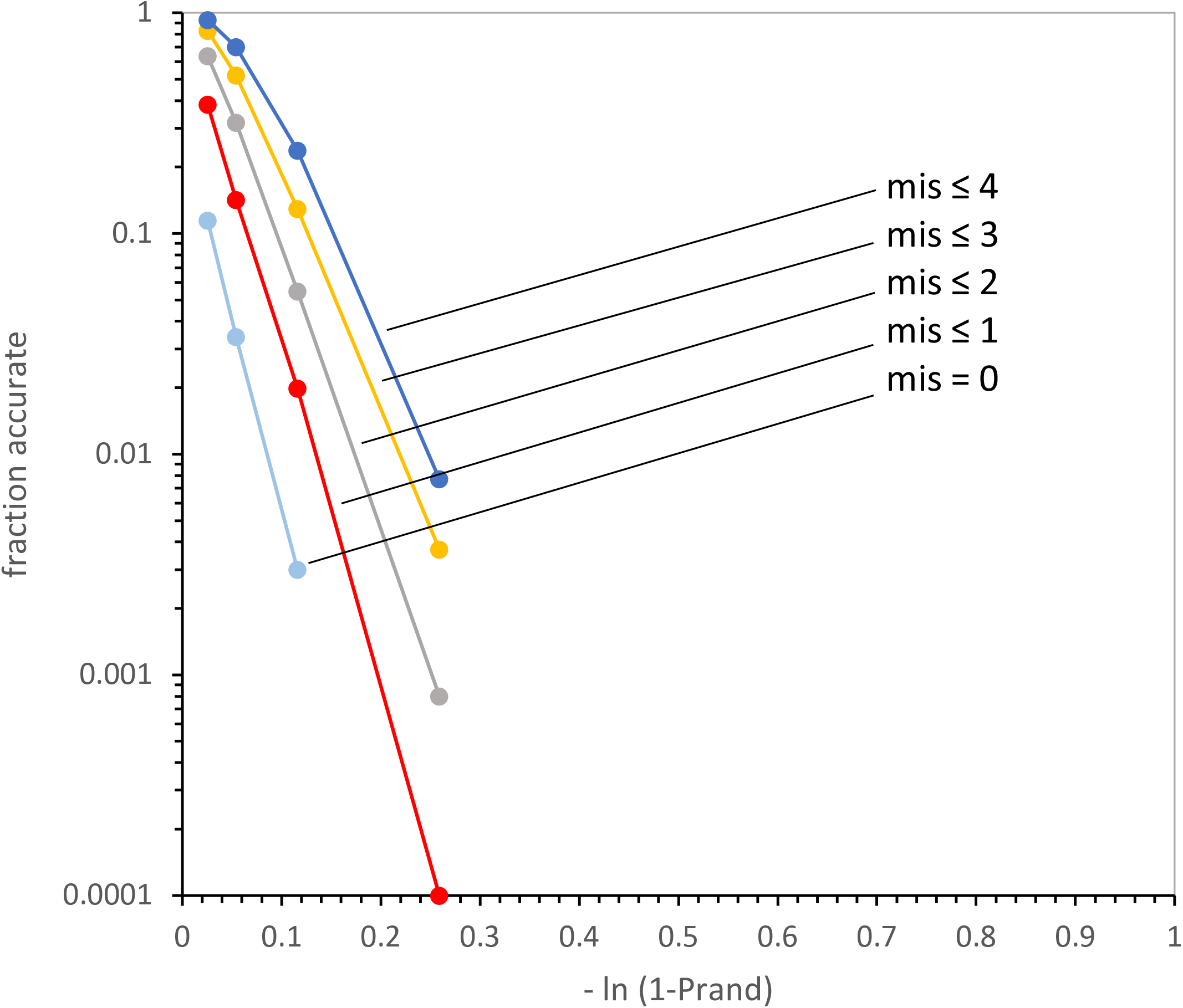
A better portrayal: a minimally evolved code and randomness: late Crick wobble, no mutational capture. Evolution of codes as in Fig. 1A (with late Crick wobble), but without mutational capture of assignments. Accuracy color coding as in Fig. 1A.

Fig. 1D replots data of Fig. 1A for complete code evolution, now including mutational capture. Comparison to Fig. 1C shows that realistic evolution is yet more sensitive to randomness, yielding ≈ ten-fold fewer accurately assigned codes (Fig. 1D). Fig. 1C and 1D taken together show that this increased sensitivity is attributable to mutational capture in Fig. 1D. This is a first indication of the negative effects of hard packing for captured wobbling triplets, explained below.

**Figure 1D.**
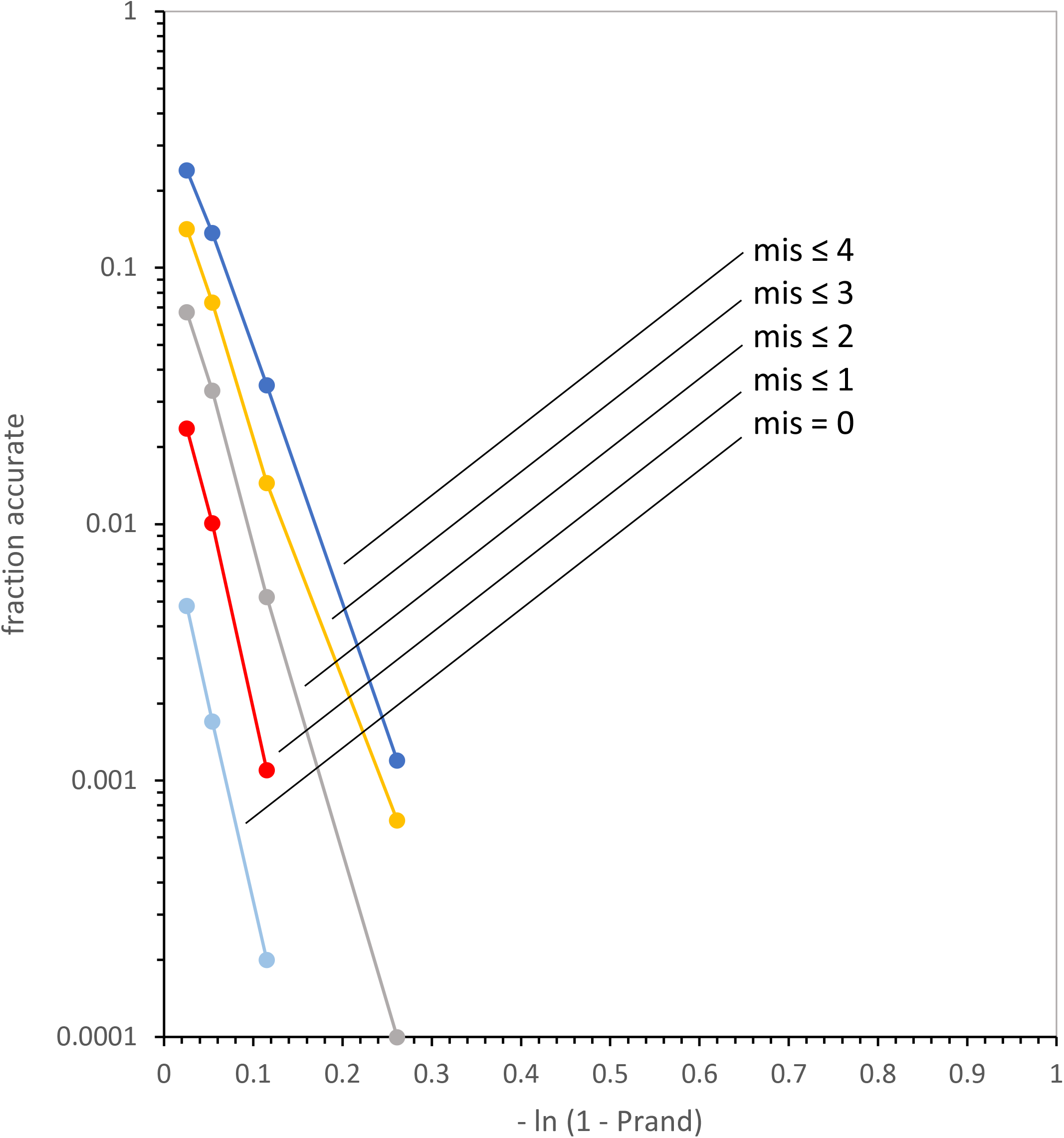
A better portrayal: sensitivity of late Crick wobble evolution to randomness. Data for complete evolution of late Crick wobble as in Fig. 1A, but plotted as in Fig. 1B. Accuracy color coding as in Fig. 1A.

### Completion complications: kinetics

Codon assignment necessarily slows near code completion, because assignment decay (which decreases assigned triplets) is speeding up, and mutational capture and triplet assignment (which increase assigned triplets) are slowing down (Yarus 2021b). Slowed late assignments show why definitive initiation and termination mechanisms were assigned late, by an independent selection, as suggested by their mechanistic differences between life’s domains (Yarus 2021c).

Fig. 2 shows that slowed code completion is intrinsic, as expected from its origin in assignment kinetics. Fig. 2A is the behavior of a minimally-evolved code as it approaches complete assignment (17 to 22 assigned functions). The figure, showing time in passages (Yarus 2021b), plots requirements for each level of assigned coding function. And, as an indicator of the efficiency of the assignment process, the mean number of assignment decays for a triplet to acquire its final meaning (decays/assignment) is also shown. Fig. 2A’s minimally-evolved coding tables are filled randomly, but with no wobble or mutational capture. Such coding still slows near completion, with 36-fold as much time required for 22 functions as for 20, requiring 53-fold as many decays per assignment for 22 encoded. For such random filling with 10% random assignments, triplets must be multiply assigned (3.5 times on average) to reach complete 22-function coding.

**Figure 2A.**
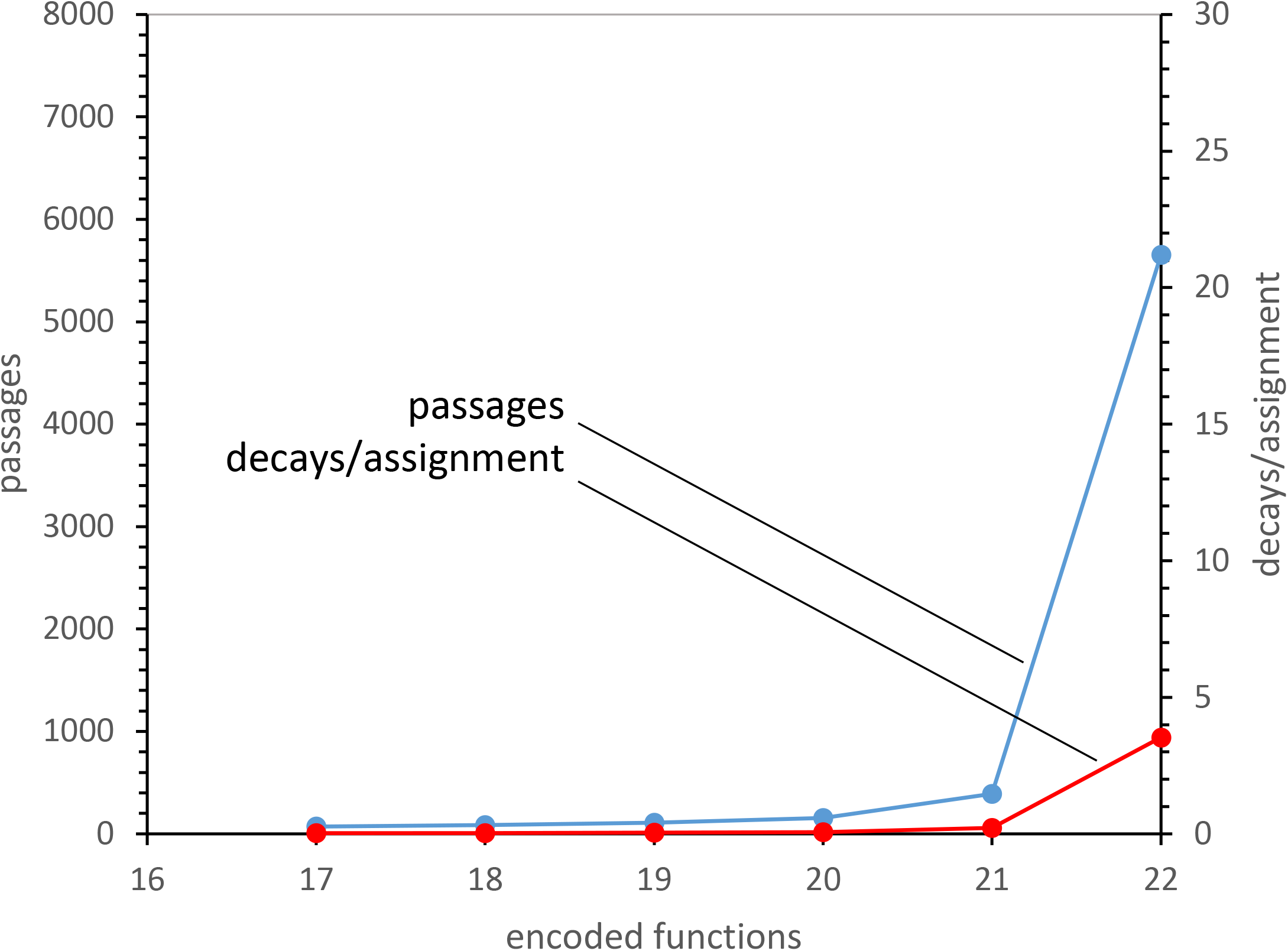
Completion complications: within a minimally evolved code. Means for 103 **c**oding tables constructed with Prand = 0.1 and no wobble, no mutational capture. Passages are computer visits to an evolving coding table, proportional to time (Yarus 2021b).

Completion becomes much more burdensome with continuous Crick wobble, even without mutational capture, as in Fig. 2B. Wobble from the start of code evolution increases both 20 to 22 function time and complexity by about 7-fold. In Fig. 2B the average triplet has decayed 24.2 times in order to encode 22 functions.

**Figure 2B.**
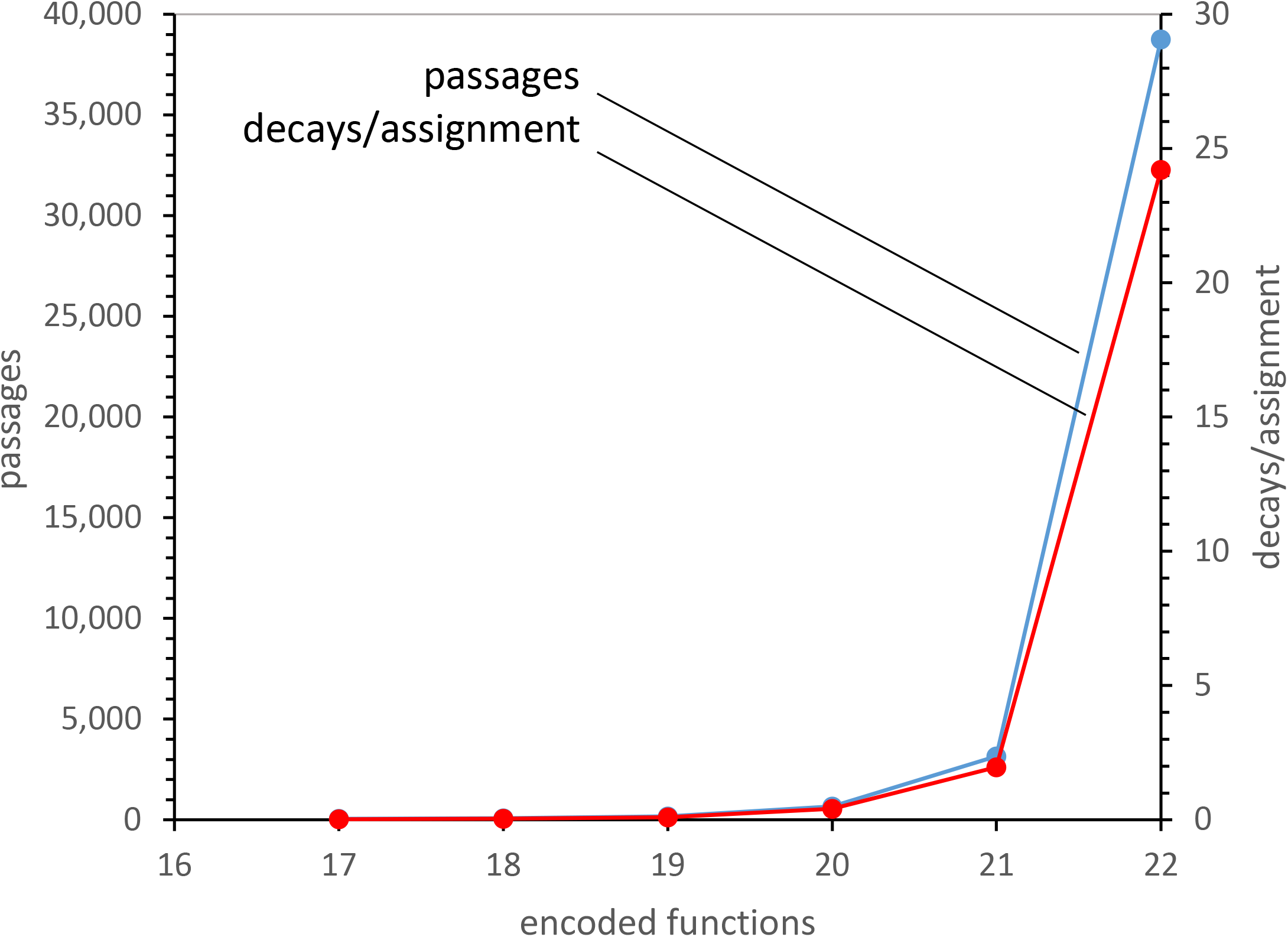
Completion complications: with wobble. Means for 10^4^ **c**oding tables evolved with Prand = 0.1 and continuous Crick wobble (that is, throughout evolution), no mutational capture.

If mutational capture of neighboring triplets for related assignments is added to Fig. 2B, as in Fig. 2C, then code completion via continuous Crick wobble is yet more hindered. To encode 22 functions, assigned triplets have decayed an average of 234 times, more than 100-fold exceeding that at 20 functions. And, time to evolve to 22 from 20 functions is > 100-fold the time to 20 encoded. Assignment of these latter triplet meanings would be tortuous, each decaying many times before 22 functions are attained. These results reproduce and extend previous comparisons in which early wobble, and early wobble to an extended range of triplets, resulted in delay and inefficient evolution (Yarus 2021b, 2021a). Increased effort for completion - from near-random filling (Fig. 2A) to continuous wobbling alone (Fig. 2B) and increased further for continuous wobbling in captured codons (Fig. 2C) further exemplify increasing difficulty in packing coding assignments.

**Figure 2C.**
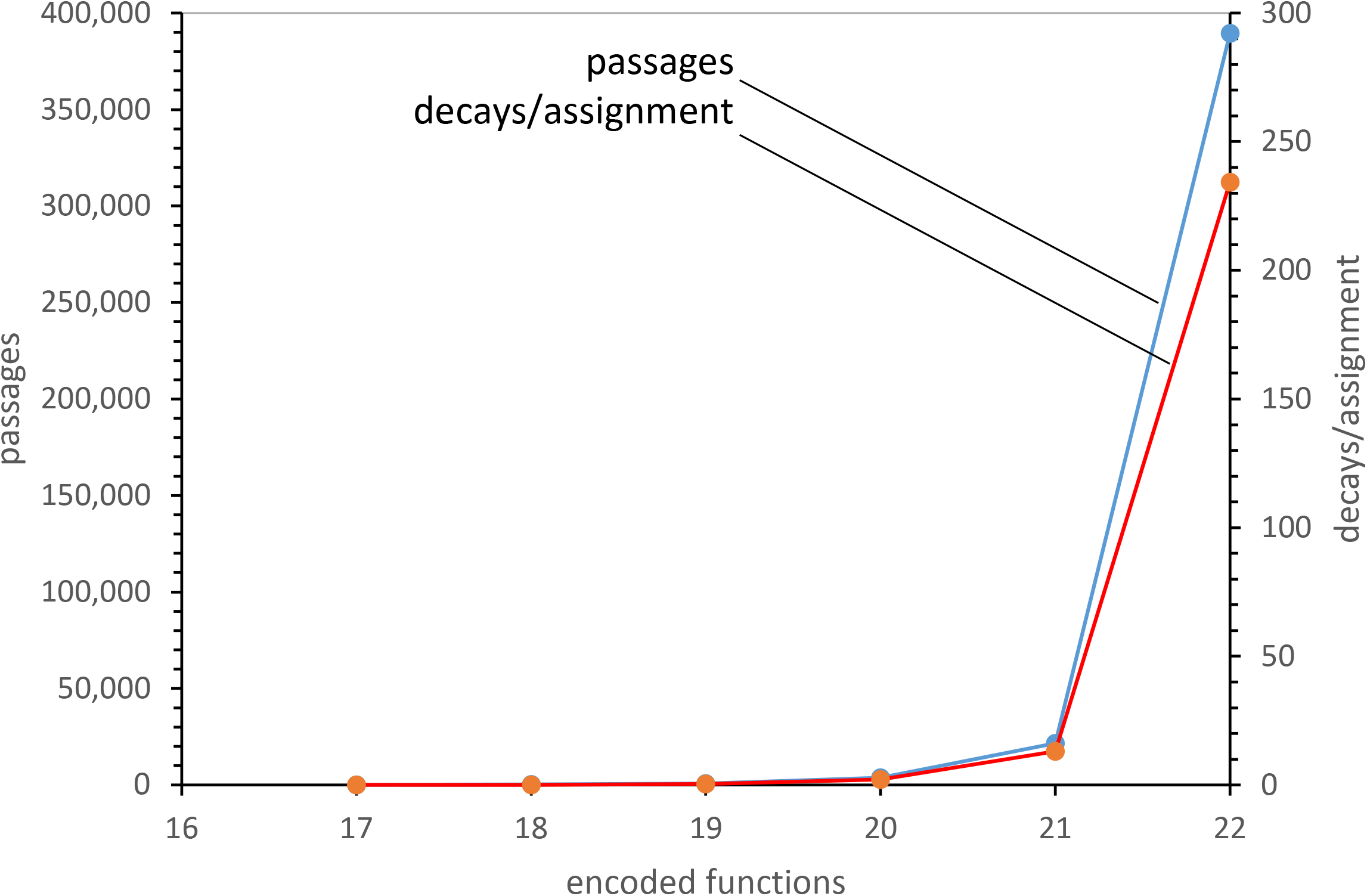
Completion complications: with wobble and mutational capture. Means for 10^3^ coding tables evolved with Prand = 0.1 and continuous Crick wobble, with mutational capture.

### Completion complications: packing

However, packing difficulties continue even if encoding 22 functions is avoided by selecting the last two functions later. In the original discussion of completion complications (Yarus 2021b), difficulties are said to be both kinetic, and due to a large universe of possible codes. Now we turn to the second kind of completion complexity, describing packing effects during the first 20 encodings.

### Hard packing: encodings that cannot overlap

Packing implies difficulty fitting code domains, like wobbling sets of codons that must preserve unambiguous meanings, into a finite, fixed coding space. Packed encodings are of differing size and complexity. In this work (Fig. 3A), triplet assignments can be unique (red), Crick wobbling (yellow, in an example initiated at AAU), superwobbling (orange, in an example initiated at GAC) or the size of the complete mutational neighborhood for capture by single mutation (blue, for Crick wobbling initiated at UUG as an illustration: Fig. 3A).

**Figure 3A.**
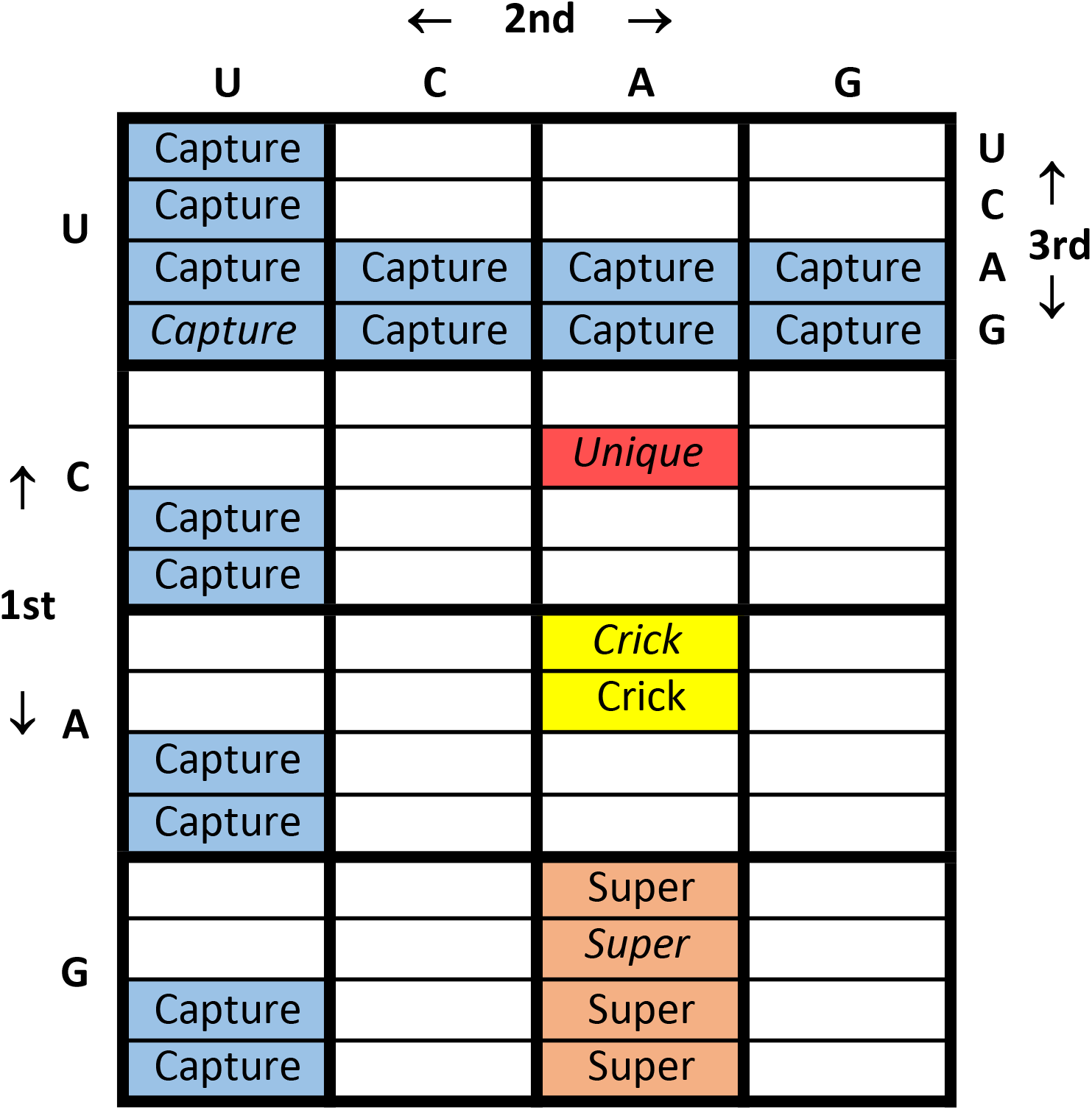
Mutational capture domains: examples in coding table context. Unique coding assigns only the target triplet. Crick wobble assigns two NNR or NNY triplets, where R = purine and Y = pyrimidine. Super = superwobble (Alkatib et al. 2012), which assigns 4 triplets, NN U/C/A/G. Capture = all codons related to example codon UUG by single mutation, followed by Crick wobble.

Pmut is the probability, per passage per neighboring triplet pair, that a chosen assigned codon will confer its meaning or a related meaning on an unassigned triplet one mutation distant (Methods). As Pmut increases for late Crick wobble in Fig. 3B, related assignments can increasingly spread across areas like Fig. 3A’s blue areas, which define a mutational neighborhood for Crick wobble initiated at the italicized *UUG* codon. Thus, as Pmut varies 40-fold in Fig. 3B, such mutational capture/related assignment goes from rare to major evolutionary event, the latter being usual in these calculations (Yarus 2021b).

**Figure 3B.**
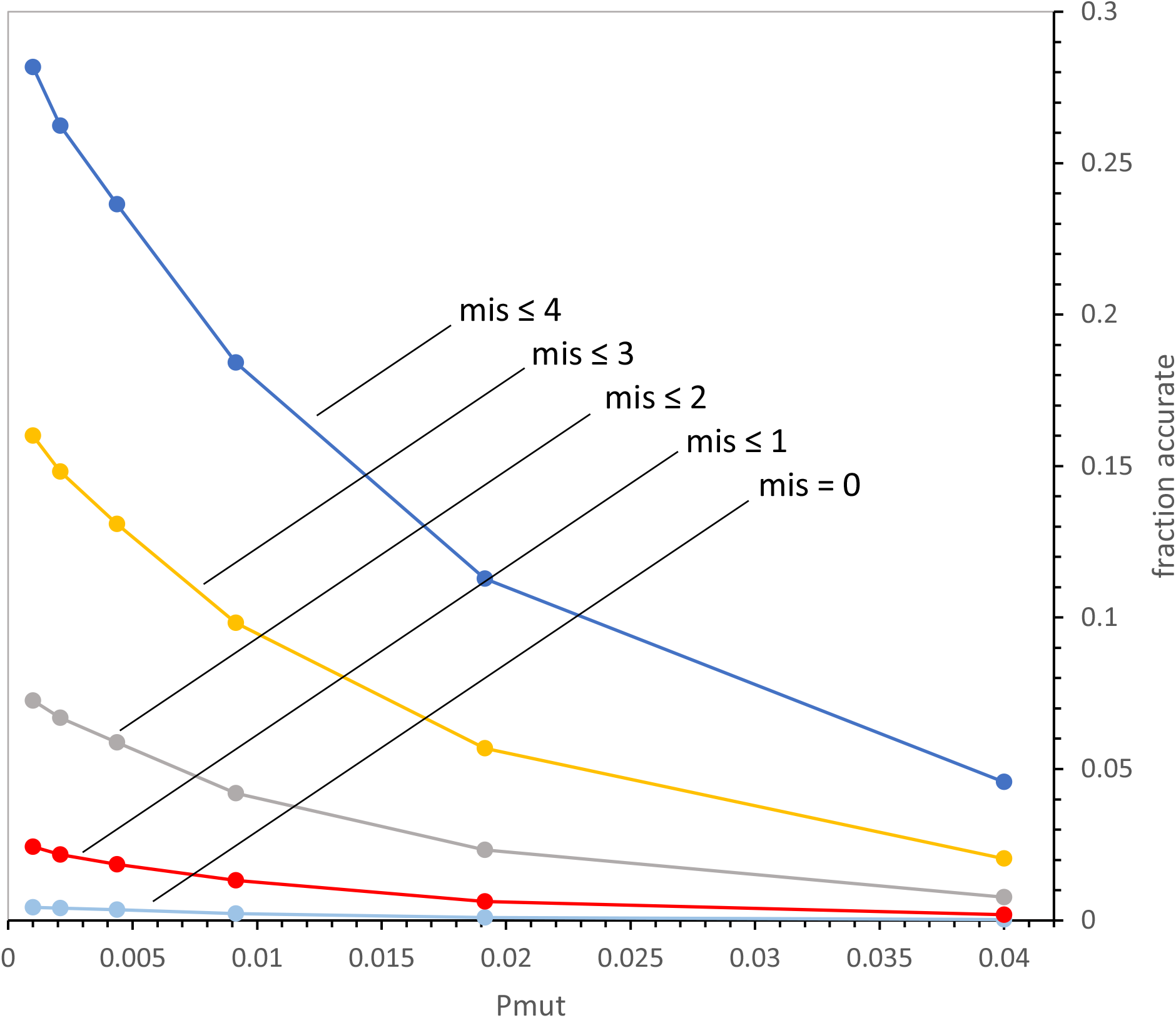
Effect of mutational capture with Crick wobble on assignment accuracy. Pmut = probability, per passage, per eligible triplet pair, for mutational capture with Crick wobble; data are means of 105 evolutions. Pmut varied from 0.001 to 0.04. Other probabilities are as in Methods. Accuracy color coding as in Fig. 1A.

Fig. 3B shows that packing is a substantial quantitative consideration. It plots probability of accurate assignment. Mean levels of codon mis-assignment relative to the SGC are plotted for 10^5^ evolved codes, varying as related assignment for neighborhood codons varies. Data is collected after assignment of 20 functions, for reasons described above, and because 20 functions are acquired near the time when best near-SGC order occurs, and therefore near the time when the SGC itself was probably selected (Yarus 2021c).

### Negative effect of assignment to a neighborhood

Mutational capture always decreases the mean accuracy of coding. Codes with ≤ 4 differences from SGC coding decrease > 6-fold with mutational capture increases of 40-fold (Fig. 2B). The negative capture effect becomes more important for greater accuracy; codes that completely emulate SGC assignment (mis = 0) are 14-fold less frequent for the same increased capture.

Fig. 3C generalizes these findings to the complete range of packings depicted in Fig. 3A. Mean fold-decreases in accuracy, measured by changed misassignment levels in 10^5^ evolutions to 20 encoded functions, are shown for the introduction of Crick wobble and superwobble into uniquely assigned codes, with (capture with the usual Pmut = 0.04, Methods) and without (Pmut = 0.001) assignment of related encodings to mutationally related codons (Fig. 3A).

**Figure 3C.**
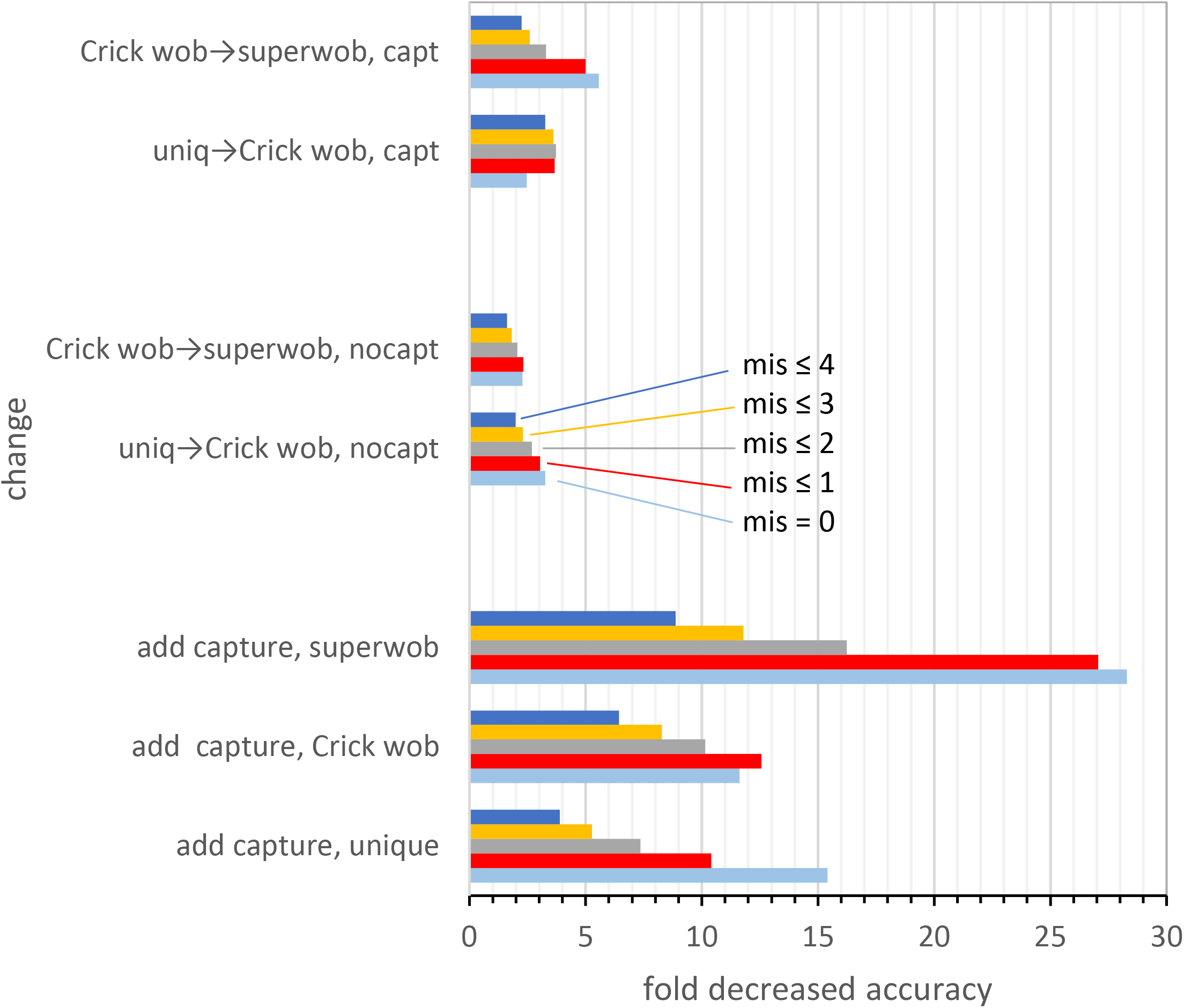
Fold accuracy decrease due to expansion of wobble capture domain: ± capture, with unique assignment, Crick wobble, superwobble. Mean effect, for 10^5^ evolutions, of the “change” described in titles on the left. Change description titles indicate, first, the conditions being compared, in a minimally-evolved background, then other relevant but constant conditions are cited. For example, “uniq→Crick wob, capt” plots the change in accuracy observed when Crick wobble is added to a minimally-evolved code that used unique assignment, both populations using mutational capture. Accuracy color coding as in Fig. 1A.

### Decrease in accuracy increases when more precision is demanded

Bundles of data in Fig. 3C present similar data for accuracy, from 4 misassignments (dark blue) at the top of each bundle of columns, to complete SGC verisimilitude (0 misassignments, light blue) at bundle bottom. A glance at the figure reveals greater decreases in accuracy as accuracy itself increases. Decreases are larger at mis = 0 than mis ≤ 4. Thus the trend in Fig. 3B is more general; packing becomes more difficult as SGC resemblance increases or misassignment decreases.

### Decrease in accuracy increases with size of the domain placed

If unique assignment is changed to Crick wobble, or similarly, Crick wobble is changed to superwobble, the size of potential wobble domains doubles (Fig. 3A); 1 to 2 triplets and 2 to 4 triplets, respectively. At the top of Fig. 3C, these similar wobble expansions have 2 to 5-fold effects on misassignment, whether their expanded wobbles are superposed on a system without mutational capture (middle data bundles) or a system with mutational capture (top bundles). On the other hand, in codes evolving with capture, which distribute their wobbles over a larger domain (top), accuracy effects are larger than when a mutational capture domain (blue, Fig. 3A) is not engaged (middle bundles, Fig. 3C). Disruptive effects ranging up to 27-fold occur for introduction of large capture domains (blue, Fig. 3A). Negative effects on assignment accuracy in these large domains clearly increase with wobble domain size: capture of unique codons is less destructive of accuracy than capture with Crick wobble; Crick wobble is less obstructive than capture with superwobble (lower three bundles, Fig. 3C).

### Hard packing summary

Accuracy penalties for packing wobble domains intensify as domains grow, and also as more accuracy is demanded. Both effects make intuitive sense, and also agree with and generalize earlier results. Previously, among populations of evolving late Crick wobble codes, the minority of codes that most resembled the SGC had also made fewer mutational captures (Yarus 2021b). Specific comparisons of unique and Crick wobble (Yarus 2021b) and Crick and superwobble (Yarus 2021a) previously favored the smaller, simpler forms. Moreover, these results (Fig. 3C, top) recapitulate the previous numerical superiority of Crick over superwobble (Yarus 2021a) - now recognizable as one sign of a more general packing effect (Fig. 3C).

What evolutionary routes minimize Fig. 3’s negative packing effects, making SGC-like assignments more accessible? Framing this as a ‘packing problem’ immediately suggests a solution: fragments of complementary shape might pack smoothly. In Fig. 4, well-known code sub-structures unexpectedly exhibit this unifying packing property. Accordingly, primordial ordered code fragments can join softly to minimize, or even entirely eliminate wobble impacts on packing.

### Primordial ordered code fragments: a row of early amino acids

(Eigen et al. 1981) noted that the most prominent amino acids from sparked gases designed to emulate primitive reducing atmospheres (Miller 1953), also have a unique coding position. These amino acids: Val, Ala, Asp, Glu and Gly, are the present occupants of the GNN row of the SGC. These findings therefore lend themselves to theories, including Eigen’s, that a primordial code would have encoded these chemically ‘primitive’ amino acids within the corresponding row of the coding table, using first-codon-position G somewhat as shown in Fig. 4A. Extensive further work on chemical properties, such as the free energy cost of synthesis in seawater (Higgs and Pudritz 2009), strikingly confirm that these five amino acids could be prevalent before biosynthesis. This grouping was also strengthened by Taylor and Coates (Taylor and Coates 1989), who noted that the same amino acids and their codons could also be classed as early, or sometimes, precisely the earliest amino acids produced from major glycolytic and citric acid cycle intermediates (as for Ala, Val, Asp and Glu). Thus chemical and biosynthetic indications concur that Val, Ala, Asp, Glu and Gly may have been early assignments to their present GNN code row (Higgs 2009).

**Figure 4A.**
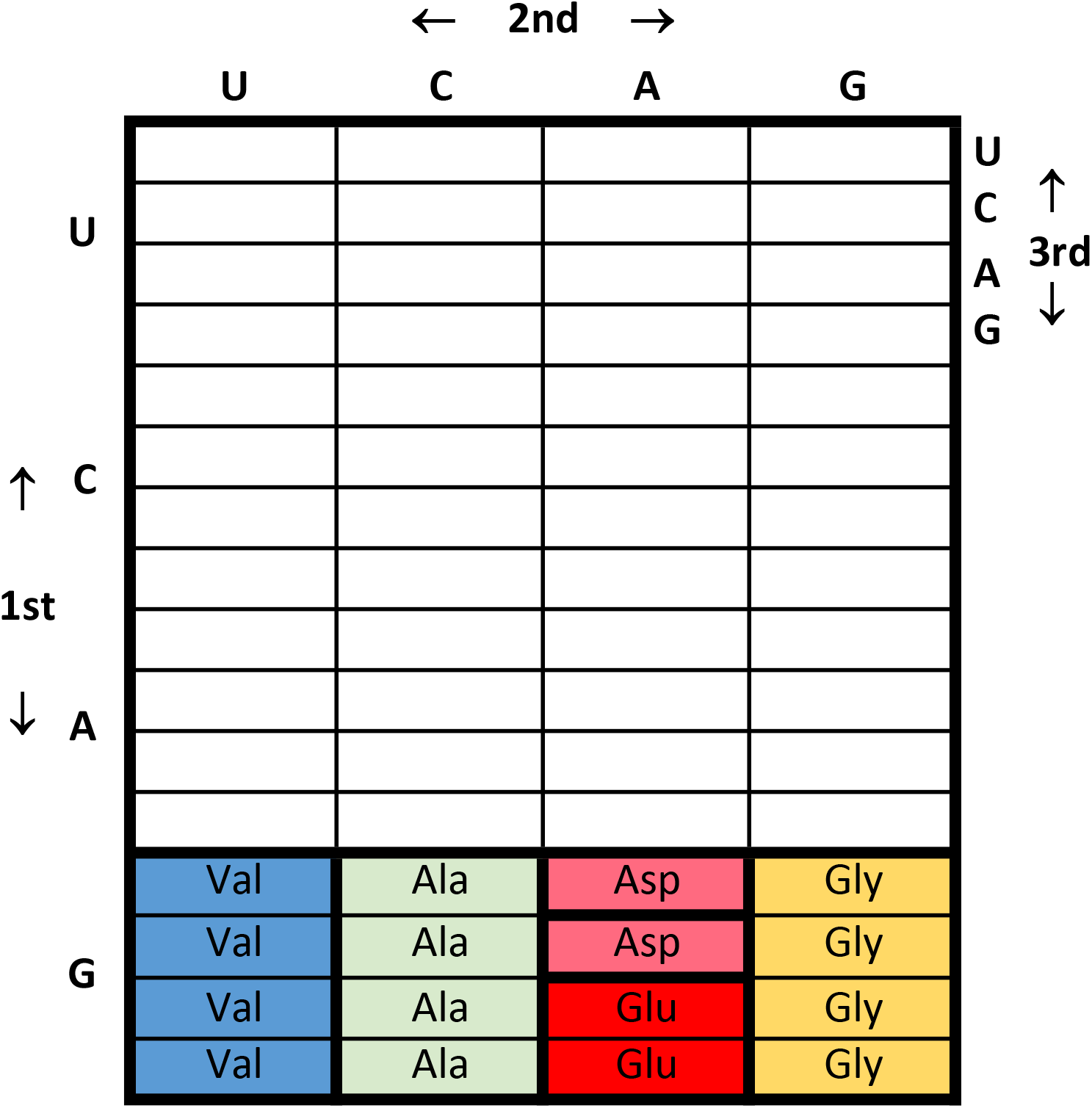
Elements of ordered coding: natural amino acids. Coding table for amino acids arguably available from ancient chemical sources, colors are arbitrary.

### Primordial ordered code fragments: synthesis and rows

A correlation between synthesis and SGC rows can be extended from the above five amino acids, arguably abundant on the early Earth, to at least 16 of 20 amino acids ultimately encoded by the SGC. Fig. 4B shows that biosynthetic origin and row coding are related for presumably later-arising amino acids, which required evolution of a biosynthetic pathway. First, in Fig. 4B, cell colors indicate likely origins from a basic metabolic intermediate (Taylor and Coates 1989): green for derivation from glycolytic phosphoglycerate, blue from glycolytic phosphoenolpyruvate, yellow from citric acid cycle oxaloacetate, and pink from citric acid cycle α-ketoglutarate. Different anabolic origins clearly tend to segregate into SGC rows, though a row relation is not completely observed. A more comprehensive summary would include minority of precursor-product relations that are related by first codon position (column) mutation (Fig. 4B). Nevertheless, frequent ordering by row suggests that the code was formed during the period when biosynthesis itself was also being established. In addition, biosynthesis approximately conserved the tendency initiated by early availability: amino acids encoded in one chemical or biochemical era tend to find assignments within an SGC row.

**Figure 4B.**
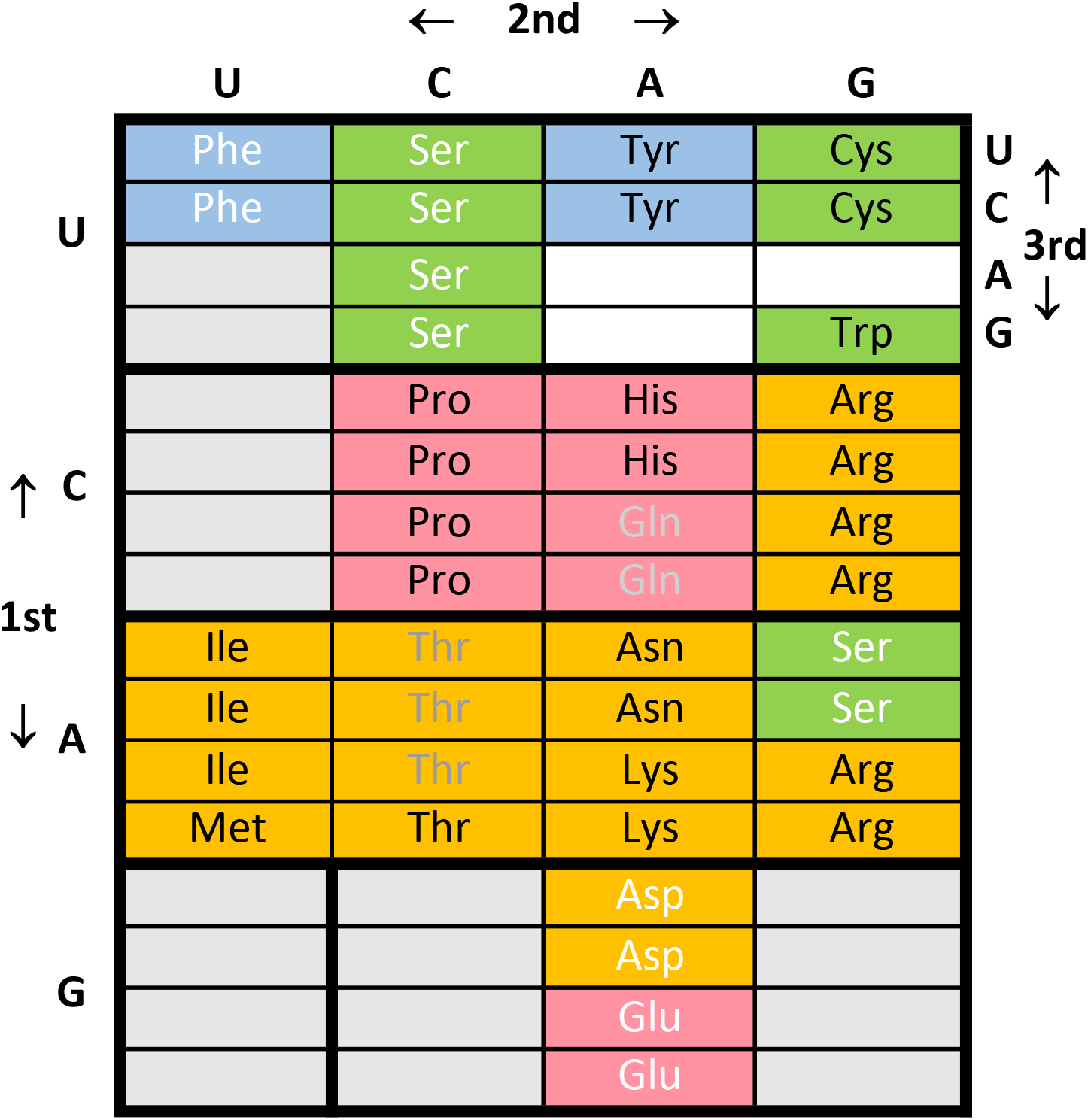
Elements of ordered coding: biosynthetic groups. Coding table for amino acids derived from glycolytic phosphoglycerate, green: from glycolytic phophoenolpyruvate, blue: from citric acid cycle oxaloacetic acid, yellow: from citric acid cycle α-ketoglutarate, pink (Taylor and Coates 1989). Text colors indicate order within a amino acid biosynthetic pathway: white, 1^st^; gray, 2^nd^; black, last.

Moreover, in Fig. 4B, text shading of amino acid names roughly indicates the order of synthesis of the amino acids: white → gray → black. Thus white Asp (from oxaloacetate) is the precursor of gray Thr which in turn is a precursor to black Ile (Wong 1975; Taylor and Coates 1989). Pooling such relations, Fig. 4B spans examples in columns and also rows. Fig. 4B’s rows and columns taken together thereby exhibit the basis for the co-evolution theory of SGC origins, in which the extension of early biosynthesis results in the assignment of closely related triplets (requiring only single mutations) to encode newly synthesized amino acids (Wong 1975; Amirnovin 1997; Ronneberg et al. 2000). Notably, such triplet concessions are not limited to rows, but have occured in the first codon nucleotide (e.g., Arg), the second nucleotide (e.g., Tyr, from Phe), and the third nucleotide (many examples, e.g., Ser, Pro and Thr).

Likely ancient chemistry and biosynthetic pathways can be fit together smoothly: (Di Giulio 2008) noted that the co-evolution of assignments and amino acid synthesis (Wong 1975) can grow from initial assignment based on ancient availability of the GNN row: Val, Ala, Asp, Glu and Gly (Fig. 4B).

### Primordial ordered code fragments: columns and amino acid chemistry

(Woese et al. 1966) suggested that the code might have originated as columns, particularly with hydrophobic amino acids partitioned into SGC columns with pyrimidines in the second codon position. Analysis of amino acid chemistry in the four coding columns (Higgs 2009) agrees, though the third (NAN) and fourth (NGN) columns of the SGC are less readily rationalized than the first (NUN) and second NCN) columns. An ancient triplet code in which the second position is the initially meaningful one is suggested by (Massey 2006); the result is a “2-1-3” model in which the SGC develops column by column, starting with the second column, then first, then third column. These ideas are approximated in Fig. 4C, which shows the SGC displayed by chemical character, using polar requirement (Woese 1965; Mathew and Luthey-Schulten 2008) as the display index. Each color represents one unit in PR, with blue the most hydrophobic (PR = 4.01 to 5.0), then light blue, gray, yellow, orange, and red the most hydrophilic amino acid (PR = 13.01 to 14.0). The SGC is plainly composed of large areas with very similar PR, amino acids whose PR is within one unit. There is a strong tendency to columns that differ, but that tend to have internally similar chemical character. A column’s tendency is not always to equality. While the third column varies continuously in PR, its structure arguably follows a continuous gradient of amino acid chemical character (Yarus 2021c).

**Figure 4C.**
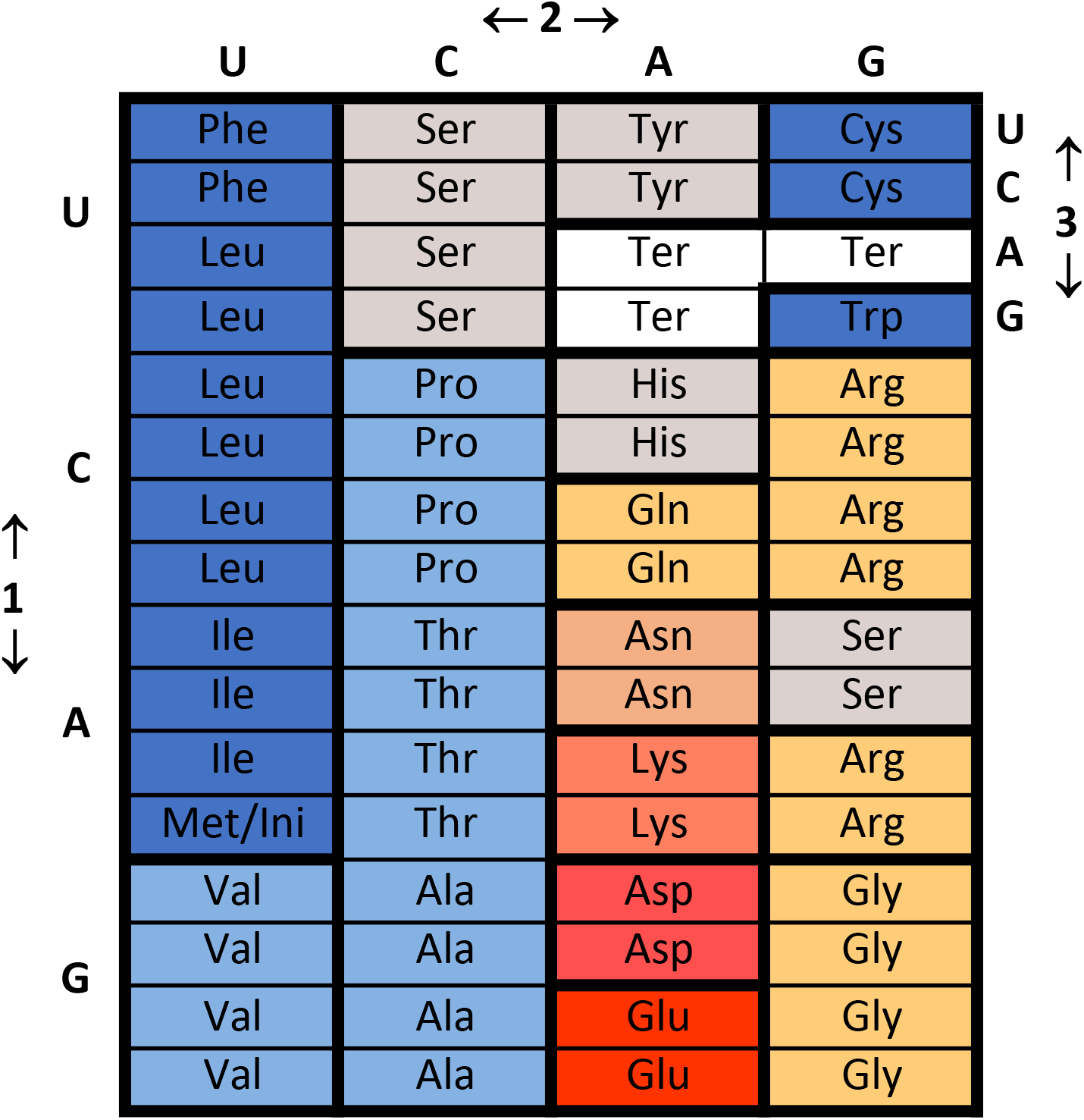
Elements of ordered coding: chemical groups. Coding table colored for hydrophobic chemical character, grouping amino acids within 1 polar requirement unit (Woese et al. 1966). From hydrophobic to hydrophilic: dark blue, light blue, gray, yellow, orange-yellow, red, deep red.

### Primordial ordered code fragments: wobble domains

It would be overlooking a major kind of SGC order to neglect the abundantly ordered wobble position, however familiar (Fig. 4D). Except for the assignments of Met/Ini and Trp, wobble ordering is universal in the SGC. The most likely explanation of SGC evolution (the Bayesian Convergence; Yarus et al. 2005) would simultaneously account for all coding regularities. Chemical character (PR) should often be conserved in columns (Fig. 4C), chemical (Fig. 4A) and biochemical origins together should tend to follow rows (Fig. 4B), and wobble behavior should almost invariably capture 3^rd^ nucleotide domains (Fig. 4D).

**Figure 4D.**
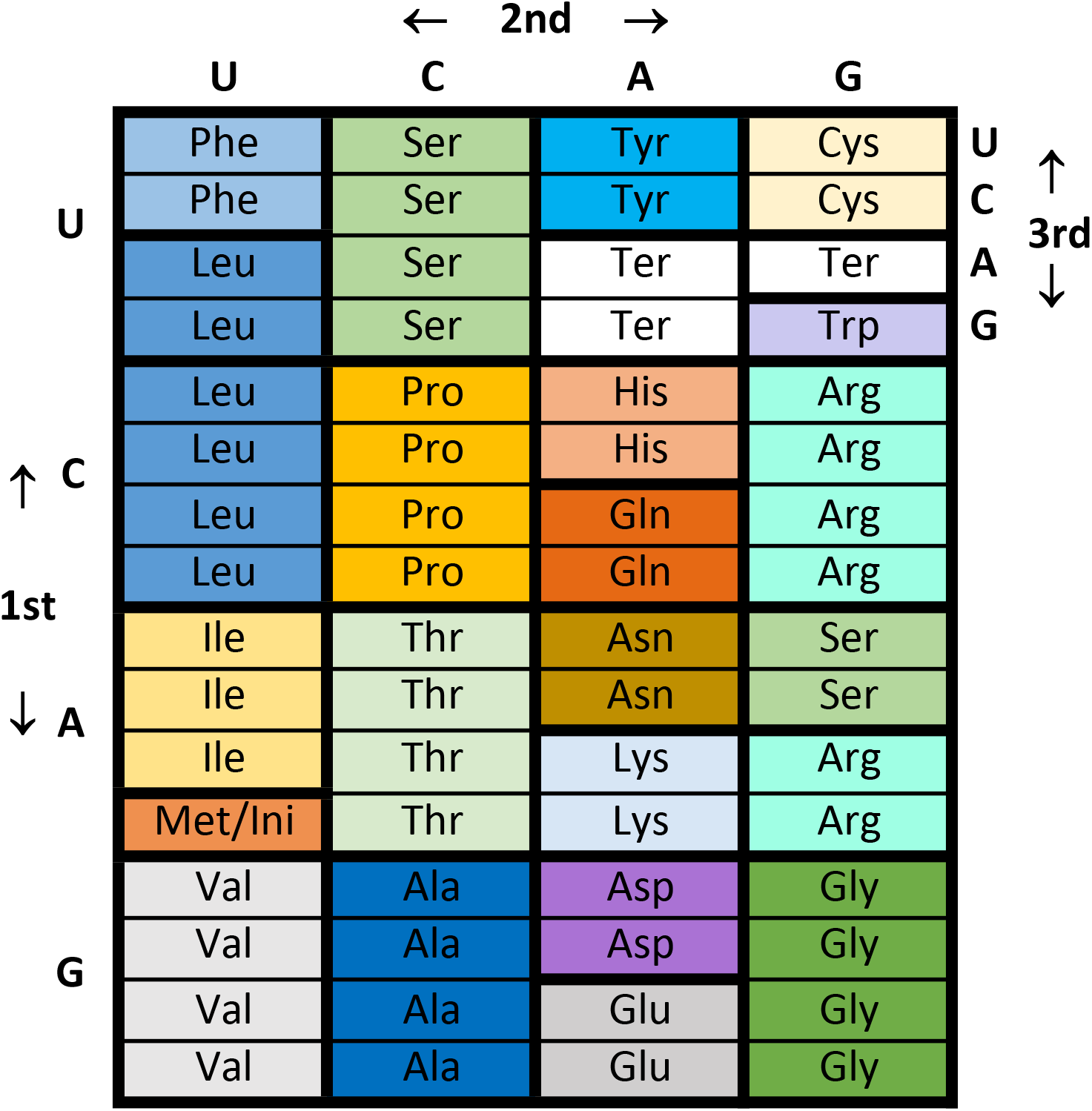
Elements of ordered coding: wobble groups. Coding table colored similarly for identical amino acids whose triplets are linked by wobble, that is, by 3^rd^ codon nucleotide variation. Individual colors are arbitrary.

## Discussion

### Non-randomness and stereochemical effects

Initial specific codon assignments are often treated as synonymous with assignments based on a definite chemical relation, like binding, between RNAs containing coding triplet sequences and amino acids (Yarus 2017). Such specific relations are called ‘stereochemical’. In Fig. 1, overall randomness in underlying assignments must be limited to <≈ 10%, in order to observe codes with 20 assignments that are also SGC-like in their encoding (Yarus 2021b). Sensitivity to randomness (Fig. 1) grows from inevitable combinatorics (Yarus 2021b). Any highly ordered code, realized via any evolutionary pathway whatever, has reached an improbable destination (Yarus 2021b). Rare SGC-like outcomes require explicit justification.

### Stereochemical origins for an SGC

On one hand, side chain- and stereo-specific RNA binding sites for amino acids have long been known (Yarus 1988), natural RNA binding sites exist (Yarus 1988; Breaker et al. 2017), bioinformatic evidence for amino acid-codon relations is plentiful (Bartonek and Zagrovic 2017), and a group of amino acid-cognate RNA sites has been experimentally selected (Yarus et al. 2005, 2009; Yarus 2017). But while cognate coding triplets recur in amino acid binding sites unexpectedly frequently, they are still sparse. Selection experiments on 8 amino acids potentially encoded by 25 codons/25 anticodons find 9 cases in which the cognate coding triplet nucleotides are improbably frequent in RNA binding sites, and are essential functional elements for amino acid affinity (Yarus 2017). Such coding triplets are especially prominent in the simplest, and therefore likely the most primitive, specific binding sites (Lozupone et al. 2003; Yarus 2017). In total, 7 anticodons and 2 codons have been detected in side chain-specific oligoribonucleotide binding sites. Thus, 28% of tested cognate anticodons and 8% of tested cognate codons appear as essential functional sequences in RNA binding sites for their cognate amino acid.

Such sparseness is unsurprising: detailed molecular interactions required for an indispensable code triplet role within specific RNA-bound-amino-acid domains would not be expected for any but a minority of tertiary structures. So, a stereochemical code origin must explain: even given multiple experimental cases of functional coding triplets, how did >≈ 90% of 61 triplet-amino acid assignments become ordered?

#### A credible stereochemical solution

The SGC can assemble from multiple smaller domains, each with non-random structures deriving from one, or a few, founding stereochemical interactions. A plausible path to the SGC then requires a way to merge these elements.

### A different SGC presentation

A more realistic visualization of codon-amino acid relationships is useful. The familiar flat rectangle compresses a three-dimensional relation (for three triplet nucleotides) to 2 dimensions. Explicitly recognizing the third dimension (Fig. 5) appreciably changes the code’s appearance, and the added dimension clarifies a complex origin (Yarus 2021b). In Fig. 5, the first two codon nucleotides extend, U to C to A to G, frontward (1^st^ nt) and left to right (2^nd^ nt). This creates sheets with the wobble nucleotide (3^rd^ nt) varying in UCAG order vertically, spanning stacked 1^st^/2^nd^ nucleotide planes.

**Figure 5.**
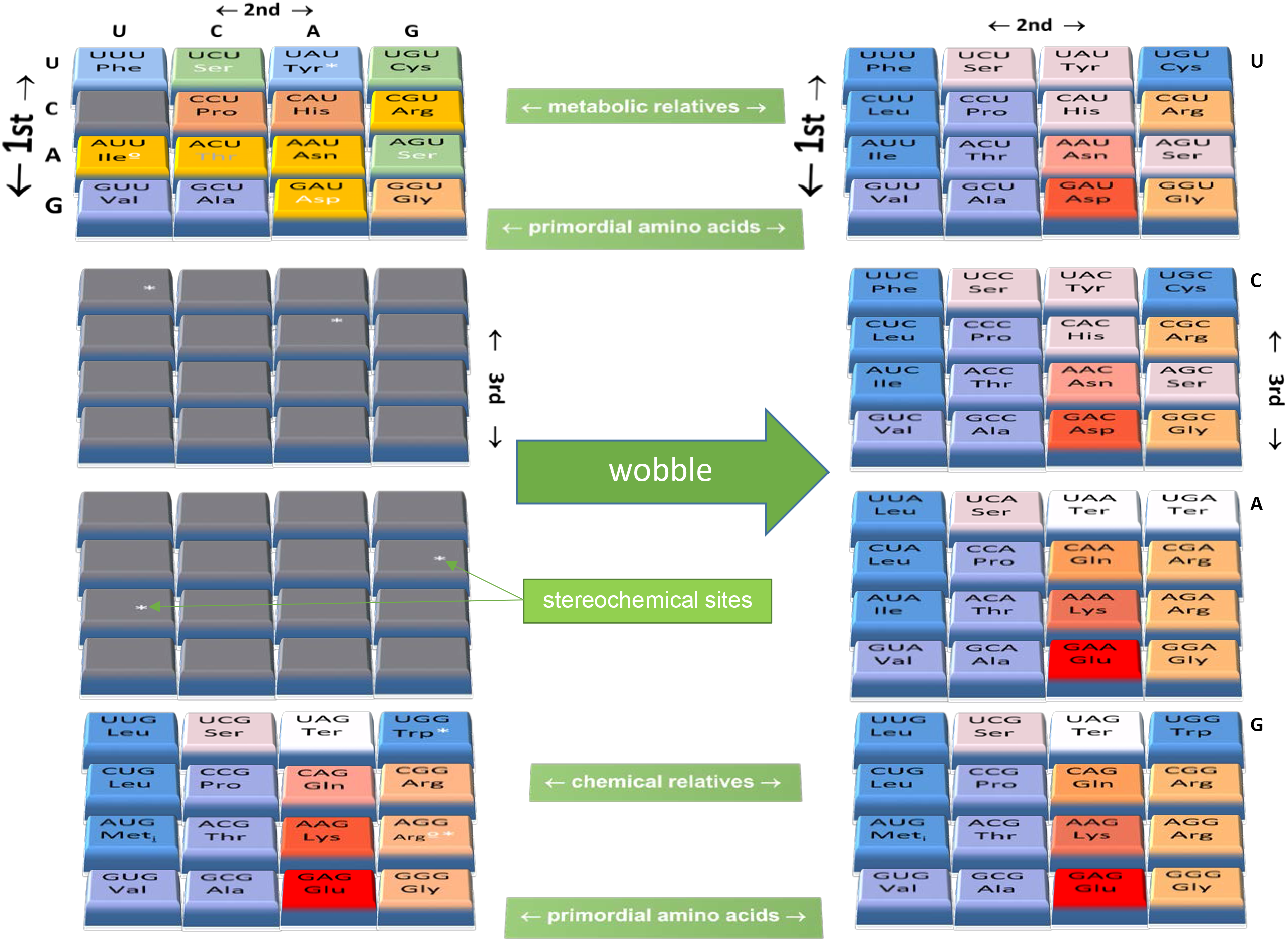
A unified SGC from three-dimensional late Crick wobble. Three-dimensional coding tables with standard assignments: 1^st^ nucleotide variation UCAG back to front, 2^nd^ nucleotide variation left to right, and 3^rd^ nucleotide variation top plane to bottom plane in UCAG order. Colors on triplet tiles correspond to those in panels of Fig. 4, explained in the text. Usual three letter abbreviations for the amino acids identify each codon assignment. Small white symbols are coding triplets exceptionally concentrated in cognate selected RNA-amino acid binding sites (Yarus 2017). Anticodon concentrations are marked with white *, codon concentrations are marked with white º. The 3-dimensional coding table on the left is a possible non-wobble, uniquely-assigned, precursor of the complete SGC on the right, which depicts coding after late Crick wobble has been adopted.

Color selection in Fig. 5 is one choice of several, but emphasizes evolutionary trends. On the left, colors resemble Fig. 4A, 4B and 4C to emphasize pre-existing ordered code domains. On the right, PR coloring (Fig. 4C) emphasizes coexistence of final SGC PR order with distinct columnar wobble domains (Fig. 4D, 5).

### The evolution shown is one variant

Fig. 5 describes the transition from an early era of unique codon assignment, before wobble (left), to a late era when Crick wobble was near-universal (right). The transition shown is not unique; it could be diagrammed with differing configurations on the left in Fig. 5, yet have SGC-like outcomes on the right (see **..from smaller ordered parts** below).

### Both hard and soft packing can exist in the SGC

Fig. 3 calculates negative effects of hard packing on SGC-like codon assignment. Hard packing characterizes regions that cannot overlap, like wobble domains that must preserve their specific amino acid meanings (Fig. 3A). However, Fig. 5 illustrates a complementary soft mode of packing, which increases SGC order. In soft packing, ordered domains readily coexist, because triplets simultaneously serve more than one role.

To illustrate soft packing, consider AUU encoding Ile (left, Fig. 5). Ile encoding has multiple associations: the codon AUU is exceptionally frequent within Ile-binding sites (Yarus 2017). The anticodon of AUA is also exceptionally frequent. Both are also prominent in maximally probable RNA-Ile binding sites, requiring a minimal number of nucleotides (Lozupone et al. 2003). But Ile AUU and AUA are also part of an extended SGC column encoding similar hydrophobic polar requirements, implying both first position (Fig. 4C) and third codon position changes (Fig. 4D). Moreover, AUU Ile is the terminus of the oxaloacetic acid biosynthetic group (Fig. 4B). Ile assignments conserve overlapping stereochemical, chemical and biosynthetic order simultaneously, because they can pack softly.

### Less hard packing conflict may be beneficial

Hard packing, as by wobble domains, unavoidably decreases assignment accuracy, especially when superposed on capture of neighboring unassigned codons (Fig. 3B, 3C). The negative packing effect on accuracy is ≈ 10-fold for late Crick wobble and high accuracies (Fig. 3C). But capture itself is not dispensable. There are many examples of chemical resemblance between metabolically unrelated amino acids assigned to SGC codons one mutation apart, suggesting capture. For example, Phe and Leu (Fig. 3B).

Some routes to reduced hard packing are evident from these data (Fig. 3). Reducing the size of a mutational capture neighborhood reduces its accuracy penalty (Fig. 3A, 3B, 3C). Perhaps the initial definition of capture (Yarus 2021b), which supposed that an assigned triplet can capture any unassigned triplet one mutation distant, was too expansive. For example, because transition mutations are usually more frequent than transversions (Vogel 1972), transition preference could define a reduced capture domain (compare in Fig. 3A). Because there is an optimally accurate early time for late wobble advent (Yarus 2021c), one could favor historical selection of the SGC on the early side of optimal, requiring fewer captures. Certainly, one should not complicate or expand simple Crick wobbles (Yarus 2021a).

### Unique amino acid assignments readily exist before wobble

Completion of a code (adding 21^st^ and 22^nd^ functions; (Yarus 2021b)) is uniquely complex for kinetic reasons (Fig. 2A, 2B, 2C). Codon assignment necessarily slows near completion of SGC evolution. Evolution of the last two functions, on average, would take much longer and require a more complex set of events than the first 20 encoded functions (Fig. 2).

This is consistent with molecular evidence that definitive initiation and termination were encoded late, after the amino acids, subsequent to separation of life’s major domains. For example, translation initiation and termination logic differ in eukarya and bacteria (Kozak 1999; Rodnina and Wintermeyer 2009) and the protein catalysts involved, necessarily themselves products of a sophisticated translation apparatus, are of independent origin in different domains. Unlike their catalysts, codons for initation/termination are near-universal, so shared primordial start/stop mechanisms may have existed also. Nevertheless, primitive but still surviving SGC encoding is probably restricted to that for amino acids.

Difficulty with latter assignments worsens if wobble occurs from the beginning of coding (Yarus 2021b, 2021a), but such barriers can be bypassed by supposing that wobble was instituted late, after the code was substantially formed by unique, non-wobbling assignments. Late wobble is independently plausible, because complex, organized ribosomal conformational changes (Powers and Noller 1990; Ogle et al. 2001)) and a particular anticodon loop conformation (Yarus 1982; Uhlenbeck and Schrader 2018) are required for accurate wobble pairing. So, before evolution of a complex translation apparatus, standard codon-anticodon base pairing, though perhaps inaccurate, is more plausible than accurate wobble. Moreover, transacylation catalyzed by RNA readily generates a suitable, simpler aminoacyl-tRNA precursor: linear aminoacyl-ribotetramers (Chumachenko et al. 2009), whose 5’nucleotides might serve an anticodon–like function (Yarus 2011; Illangasekare and Yarus 2012; Yarus 2021b).

Thus: early coding may still be visible in unique amino acid assignments, without Crick wobble. But the present SGC likely appeared subsequent to late Crick wobble, defined as NN U/C and NN A/G translation by individual acceptor RNAs (Yarus 2021b). An earlier non-wobbling SGC precursor is implied, with near-complete amino acid assignment. Fig. 5 (left) shows that this implied non-wobbling SGC precursor can exist. Further, the non-wobbling precursor is a sufficient foundation for the complete row- and column-ordered SGC (Fig. 5, right).

### The influence of likely ancient amino acids is preserved

A broad consensus finds Val, Ala, Asp, Glu and Gly to be credible primordial amino acids for reasons of synthetic ease (Miller and Urey 1959), natural abundance (Cronin et al. 1979) and ready thermodynamic access (Higgs and Pudritz 2009). In Fig. 5, this probable primordial status is accepted: all code order ultimately rests on initial encoding of these amino acids by the row of GNN codons in the SGC.

Amino acids probably available before biosynthesis (Fig. 4A) occur in both the NNU (upper) plane and the NNG (lower) plane. The upper SGC plane fuses the primordial chemistry row (Fig. 4A) with the row-biased biosynthetic domains of Fig. 4B. The lower SGC plane combines primordial availability (Fig. 4A) with the SGC’s chemically organized columns (Fig. 4C).

### No recoding is required to add late wobble

In Fig. 5, wobble advent requires no changes whatever to initial encoding. This is notable, because wobble as a source of SGC order affects most encoded functions (Fig. 4D); its influence on code structure spans the coding table. Wobble has a major evolutionary role in shaping the SGC’s close spacing of assignments for related functions (Yarus 2021b). Moreover, late wobble is the most probable path to full and complete codes (Yarus 2021b, 2021a). It is noteworthy that a fundamental coding feature can be added after most assignments are made, without perturbing extensive foundations.

This is particularly true in light of negative wobble effects. Larger wobble domains, and earlier wobble also, increase completion complications (Fig. 2). This is more important when larger wobble domains are spread to a triplet’s mutational neighborhood by capture of nearby unassigned triplets (Fig. 3). Decreases of more than an order of magnitude in assignment accuracy are routine for use of capture and wobble, with the greatest difficulty occurring for the greatest resemblance to the SGC (Fig. 3B, 3C). These completion complications are entirely relieved via the costless pathway for late-wobble introduction shown in Fig. 5.

### The SGC is accessible from experimental levels of RNA-amino acid stereochemistry

In Fig. 5 (left) small white symbols on triplet tiles indicate conserved, functional, cognate coding triplets within experimentally-selected amino acid binding sites. White symbols indicate 8 triplets with 9 experimental stereochemical connections, quite varied in amino acid side chain and coding sequences: Ile codon AUU, Tyr anticodon AUA, Phe anticodon GAA, His anticodon GUG, Ile anticodon UAU, Arg anticodon UCG, Trp anticodon CCA, and uniquely: Arg, employing both codon AGG and its anticodon CCU (Yarus 2017).

Nevertheless: how were most SGC sense triplets ordered, beginning from ≥ 8 initial loci? Fig. 5 takes experimental RNA-amino acid interactions as nuclei for small, ordered code substructures. In this way, each left-hand SGC plane has two documented stereochemical nucleations. Thus the SGC, unexpectedly, can appear over-determined; there are so-far functionless stereochemical sites in unassigned gray areas (Fig. 5, left). Such excess suggests an additional role for amino acid-RNA interactions in onset of Crick wobble, between Fig. 5’s left and right halves.

### A stereochemical role in wobble advent

There is a straightforward way, economical of hypotheses, to add unused stereochemical assignments (in Fig. 5, left) to late wobble evolution. A two-plane model may be oversimplified, given that there are possible stereochemical triplets and likely ancient Val/Ala/Asp/Glu/Gly assignments on all four SGC planes (Fig. 4A). Thus, soft packing of ordered code precursors might occur through all planes (Fig. 5), utilizing all known amino acid-RNA relations (Yarus 2017). This will bear more thought.

### Sources of SGC order coexist

Previous analysis (Yarus 2021b) suggests that, in the very highly ordered SGC, related assignment to chemically similar (Woese et al. 1966), related assignment to biosynthetically related amino acids (Wong 1975) and to chemically similar amino acids (Freeland and Hurst 1998) should occur simultaneously. Further, chemically determined assignments and error minimization appeared to be statistically independent (Caporaso et al. 2005). It has not been clear how such differing molecular objectives could be satisfied together in the SGC. Fig. 5 now shows that different principles can be implemented in separated SGC regions (left) and later combined (right) after accurate wobble evolves.

### Selection acts at the newly defined level

The fit of ordered code parts, simultaneously preserving primordial origins (Fig. 4A), biosynthetic relations (Fig. 4B) and chemical function (Fig. 4C) while allowing wobble coding (Fig. 4D, 5) is surely not accidental. Instead, it implies a new dimension for distribution fitness, that is, evolutionary selection of extreme members of a broadly-distributed population (Yarus 2021b). Via Fig. 5’s pathway, selection for efficient translation chose, from a varied population of intermediate codes, complementing parts that merged into a complete SGC.

This is exemplified by a specific 3-dimensional late Crick wobble example. Earlier unique coding might make different assignments to NNA and NNG, or alternatively, to NNU and NNC codons. But using Crick wobble (Yarus 2021b), neither NNA nor NNC can be assigned specifically, because later wobble will make them equivalent to NNG and NNU, respectively. However, with freedom to select combined partial codes as in Fig. 5, such conflicts can be avoided; an SGC-like group can be found among an unchanged population of nascent codes.

### Late Crick wobble precisely generates full codes

Code completion can be variously defined. One might be interested in ‘full’ codes (all triplets assigned) or ‘complete’ codes (all functions encoded). Prior results (Yarus 2021b) suggest that late Crick wobble approaches full coding more precisely than continuous wobble, while still allowing coding capacity for later definitive initiation and termination. Three-dimensional late Crick wobble (Fig. 5) now makes superior access to full coding explicit. Late Crick wobble among uniquely-assigned precursors (Fig. 5, left) precisely fills the coding table, closely approaching the full 3-dimensional SGC (Fig. 5, right) via one uniform transition. This recalls the finding (Yarus 2021b) that unassigned triplets readily persist into late code history, and therefore could take late evolutionary roles.

### AN NNG intermediate appears a reduced, capable, translation system

In Fig. 5, the lower proposed plane encodes 13 amino acids, as well as early initiation and termination. This is particularly striking; a chorismate mutase has been reduced to 14, then to 9 amino acids (Walter et al. 2005). Reduction of the amino acid complement in nucleoside diphosphate kinases shows that stable structures and enzymatic activities are accessible together if 13 different amino acids are encoded (Kimura and Akanuma 2020). The NNG plane (Fig. 5, left) therefore is a particularly plausible evolutionary intermediate because it independently encodes accurately terminated, functional enzymes.

### An intricate development is facilitated by division into regions

SGC-like codes arising earlier would more probably be selected. Division of SGC encoding into multiple regions allows code evolution in parallel, which potentially shortens time to completion, compared to a single linear path.

This repeats a common theme; evolution in multiple regions is also used to minimize penalties for hard wobble packing (Fig. 3). Evolution in multiple regions allows simultaneous implementation of varied means to code order (Fig. 4, 5). Selection of multiple regions acts on smaller, perhaps more readily organized, coding intermediates (Fig. 1, 5).

Moreover, evolution by division shrinks population size required to select a code. This size is defined by the abundance equation, S = ln 2/P_SGC_ (Yarus 2021c), where S is the number of independent codes that must be examined by selection to find, with probability 0.5, a code occurring with fractional abundance P_SGC_. However: reassorting preexisting partial codes eventually allows all combinations, diversifying codes available from a fixed number of individuals, thereby shrinking the population required to select an SGC.

### The ordered SGC was assembled from smaller ordered parts

The SGC likely originates as less functional partial codes packed softly to fill a coding table (Fig. 5). Relevant evolution has been studied quantitatively (Sella and Ardell 2006). The crucial idea is that newly evolved codon assignments are constrained by history, because new assignments must preserve function of already-encoded peptides. Such conservative selection is sufficient to create coding that approaches SGC levels of order. Such order evolves, even beginning with a code that is uniformly ambiguous (Sella and Ardell 2006), so that no relation whatever between codons and amino acids preexists. Ordering works better after a few early assignments, and so is well-suited to a few prior stereochemical assignments (Fig. 5, left). It readily produces 2-dimensional codes for ≈ 12 amino acids, explicitly supporting planar intermediates like those in Fig. 5 (Sella and Ardell 2006). Such conservative evolution should therefore transmit an overall chemical logic from initial assignments (Yarus 2017) to subassemblies (compare **An NNG intermediate..** above), then to the SGC (Fig. 5, right).

### Origin of smaller ordered SGC parts

Possibly, ordered SGC subsections evolved in primitive cellular compartments, forming the SGC by compartmental membrane fusions. However, an alternative pathway emulates accepted later events in cellular evolution.

About 2.2 billion years ago, as oxygen initially accumulated in the Earth’s atmosphere (Schopf 2014), an α-proteobacterium (Gray 2012) began an endosymbiotic relationship with a still-uncertain archaeon (Archibald 2015) related to Asgard archaea (Spang et al. 2018). The bacterium became the ancestor of the eukaryotic mitochondrion, and thus of a chimeric aerobic eukaryote. Chimerism was followed by transfer of numerous genes from endosymbiont to host cell (Ku et al. 2015). Multi-compartment eukaryotic cells notably have distinct genetic codes in different compartments, sometimes relying on transferred tRNA genes (Jukes and Osawa 1990). A remarkably parallel sequence of events ≈ 1.5 billion years earlier, in which ancient cells with differing, partial codes fused, could have founded the SGC. Vetsigian (Vetsigian et al. 2006) suggest that horizontal gene transfer created a universal SGC. Horizontal gene transfer, long before universality, could also have created the ancestral code (Fig. 5).

## Methods

### Computed coding table evolutions

Simulations have been described with more detail (Yarus 2021b, 2021c). Calculations begin with an empty coding table. Code evolution is divided into short time slices. In each slice, a nascent coding table is visited by computer. A random triplet is chosen. Each such choice initiates a computed passage, during which only one event: initiation or decay or capture or alternatively, nothing at all, will occur. If the chosen triplet is unassigned, it can be assigned to one of 20 amino acids, initiation or termination. Assignments can be to one triplet (unique), or to a group related by wobble. Subsequent assignment decay also occurs uniquely, or for a wobble group, if such a group exists.

All events happen stochastically, determined by randomized numbers, conferring assigned probabilities for one passage. These procedures are equivalent to assigning chemical rate constants (Yarus 2021b), assuming that initiation and decay are first order in unassigned and assigned codons, respectively, and that mutational capture is second order, depending on the product [assigned triplets*unassigned triplets] (Yarus 2021b). An important implication is that passages are an appropriate time unit, proportional to real-world time.

It is assumed that representative probabilities for evolutionary events during one passage exist: for example, probability Pinit for initial codon assignment, Pdecay for loss of assignment, Pmut for mutational capture of an unassigned codon one mutation distant (using the coevolution (Wong 1975)/polar requirement (Freeland and Hurst 1998) protocol called Coevo_PR (Yarus 2021b)) and Pwob for wobble when there is a choice between types of wobble or unique encoding. Prand is the probability that an initial assignment is random, with no specified relation to the same SGC triplet’s meaning. Here, unless otherwise specified (for example, when a probability is being varied): Pinit = 0.6, Pdecay = 0.04, Pmut = 0.04, Pwob = 1.0, Prand = 0.1 for each passage.

Example source code comprising ≈ 900 lines of Pascal, and an Excel spreadsheet example of downstream analysis and graphics, are available at 10.5072/zenodo.733491.

### Minimally-evolved genetic codes

These have specified numbers of randomly chosen triplets with SGC assignments, complemented by completely random assignments for other codons. Such coding tables may not have mutational capture, and do not evolve in other ways. However as used here, minimally-evolved coding tables do allow prior assignments to decay. Otherwise it can be impossible to complete a coding table that has a requested composition or history.

Without decay, such a calculation can hang indefinitely, unable to recover from a poor assignment. In contrast, when decay is possible, difficult evolutions appear instead with a longer completion time, and are readily included in population statistics.

### Assignment accuracy for fully random coding

In Fig. 1B, average accuracies are plotted for coding tables filled randomly. These are calculated from the binomial distribution:

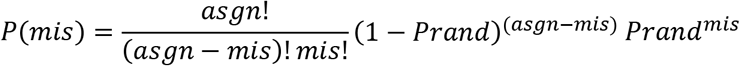

where *mis* is the number of misassigned triplets, *P(mis)* is the probability of *mis* missignments when random assignments occur with *Prand*, and *asgn* is the total number of assigned triplets. *P (mis)* for different *mis* were summed as required to get results in Fig. 1B. Fig. 1A has a mean *asgn* = 58.56; here this mean distribution is used alone to approximate distributed occupancies. Sums of the natural log of *P (mis)* should have an innate dependence on ln (1 – *Prand*), as shown by the above equation and in Figures 1B, 1C and 1D.

## Acknowledgement

Thanks to Tom Cech for discussion of a draft manuscript.

## Notes

### Competing Interest Statement

The authors have declared no competing interest.

### Summary of Updates

Numerous small clarifications have been made.

